# The immunomodulatory p43 secreted protein of *Trichuris* whipworm parasites is a lipid carrier that binds signalling lipids and precursors

**DOI:** 10.1101/2025.03.09.642242

**Authors:** Malcolm W. Kennedy, Allison J. Bancroft, Richard K. Grencis

## Abstract

Parasitic helminths such as the *Trichuris* whipworms contribute to significant disease and morbidity in humans and other animals. They cause prolonged infections despite intact immune systems of their hosts, and their persistence is increasingly attributed to immunomodulatory activities of their secreted products. We examined the p43 (Tm-DLP-1) protein of *Trichuris muris* that comprises about 95% of the protein secreted by the adult parasite, is known to bind matrix proteoglycans, and have IL-13-neutralising activity. We show here that worm-derived p43 purified from secretions binds fatty acids and retinol, including signalling lipids or precursors thereof. This binding activity might therefore contribute to the parasite’s modulation of the infection site by delivering or sequestering influential lipids. A recombinant orthologue of p43 from the human whipworm, *Trichuris trichiura* (p47; Tt-DLP-1), has similar lipid-binding activity, with a similarly highly apolar biding site. From the known molecular structure of p43, we note the existence of extensive surface-accessible cavities with diverse surface charge characteristics which may indicate binding of a range of small molecule types in addition to lipids, and that its internally duplicated subdomains likely possess discrete characteristics. p43 belongs to the “dorylipophorins” that have only been found in Dorylaimia (Clade I) nematodes and appear to be the chief lipid carrier in their pseudocoelomic fluids, replacing the major lipid transporters of other nematode clades. In *T. muris*, and potentially other trichurids, these molecules appear to have been adapted for both internal physiological and external immunomodulatory activities.

## Introduction

Parasitic nematodes, both gut- and tissue-parasitic, are remarkable in the chronicity of their infections in the face of otherwise intact immune systems of their hosts that they are able to modify to their advantage ^1–4^. Over recent years it has become increasingly apparent that their secretions are key to their immunomodulatory activities, the active principles ranging from proteins, adducts to proteins, extracellular vesicles, to lipids identical to those used by their hosts to regulate their own immune and inflammatory responses ^2,4–16^. Experimentally, purified natural or recombinant forms of each of these classes of material can be shown to be active, but it is also clear that each species of parasite likely uses a combination of effectors (see above).

Here we examined the p43 protein of *Trichuris muris*, the intestinal whipworm of mice, which is by far the most abundant protein in the secretions of the adult parasites. It has been found to bind and compromise the activity of IL-13, the crucial cytokine in the immune rejection of this parasite ^17^. Moreover, p43 binds glycosaminoglycans (mucopolysaccharides), thereby localising and tethering the protein to its presumed site of action proximal to mucosa. Remarkably, there is only a limited antibody response, and no discernible T cell response, to p43 during infection, even in chronic infections with the parasite ^17^. This may reflect its confinement largely to the intestinal niche despite its interaction with crucial components in the immune system. The anterior of *Trichuris* species is burrowed into the gut epithelium ^17,18^ of the colon, which could expose the immune system to the protein, but it is not known from where in the worm that p43 is released, or whether there is some mechanism of induction of immune tolerance. Parasites of the *Trichuris* genus such as *T. muris*, *Trichuris trichiura* of humans, and *Trichuris suis* of pigs, share similar life cycles and morphological similarities such that it seems likely that they use similar mechanisms by which to modulate the immune systems of their hosts, and that their p43 orthologues may therefore play similar roles.

We show here that both the p43 protein of *T. muris* and its orthologue (p47) of *T. trichiura* bind lipids including fatty acids that can be precursors of a wide range of signalling lipids involved in cellular interactions, including immune and inflammatory responses. Moreover, p43 is the major protein in the pseudocoelomic fluid of *T. muris*, where it probably acts as a bulk mobiliser and transporter of lipids internal to the worms as a functional equivalent of the major pseudocoelomic proteins of other clades of nematodes ^19–24^. In *Trichuris* parasites at least, p43 has been additionally devoted to a function external to the parasite. A lipid-binding protein released by a gut parasite might therefore modify local immune responses by delivering active lipids, or sequestering host signalling lipids. The indications are hence that the p43 protein of *Trichuris* whipworms is a major internal lipid transporter of a type common to Dorylaimia nematodes, but additionally adapted for immunomodulation of hosts.

## Materials and Methods

### Animals and ethics statement

Severe combined immunodeficient (SCID) mice were bred in-house at the Biological Services Facility at the University of Manchester and used to provide adult *T. muris* worms for the provision of parasite-derived materials as described. Procedures were performed under the regulations of the Home Office Scientific Procedures Act (1986) and were subject to local ethical review by the University of Manchester Animal Welfare and Ethical Review Body and followed ARRIVE guidelines for animal research.

### Nomenclature

The p43 protein of *Trichuris spp.* is an orthologue of the poly-cysteine and histidine-tailed protein (PCHTP) originally described from *Trichinella spiralis* ^25^, and the P44 pseudocoelomic fluid protein of the giant kidney worm *Dioctophyme renale* ^26^. This family of proteins appears to be confined to Clade 1 nematodes, the Dorylaimia ^27,28^, hence the proposed name of doryliphorins (DLP), given their here-defined biochemical activity and class of nematodes which have them. The systematic name of *T. muris* p43 would therefore be Tm-DLP-1, and of *Trichuris trichiura* would be Tt-DLP-1, but we use the terms p43 and p47 (based on their apparent molecular masses) here for simplicity and continuity.

### Worm-derived materials

*T. muris* p43 protein was produced from adult worms that were cultured *in vitro* to obtain their excretory/secretory (E/S) material from which purified p43 was produced, all as described in ^17^. Briefly, medium was collected from adult worms cultured for 4 hours and overnight in RPMI-1640 with 500 international units/ml (IU) penicillin and 500 μg/ml streptomycin. This was centrifuged to removed eggs and filtered through a 0.22 μm syringe filter. The medium was incubated overnight with Ni-NTA Agarose (Qiagen, Manchester, UK) at 4°C on a rotator. The p43 protein was eluted from the Ni-NTA with 250 mM imidazole and further purified using a size-exclusion column (Superdex 75; GE Healthcare, UK) and the relevant fractions were concentrated, filtered through a 0.22 μm membrane, aliquoted, and stored frozen.

Pseudocolemic fluid (PCF) was collected from adult *T. muris* by placing worms that had been rinsed in culture medium in a small amount of RPMI with 500 U/ml penicillin and 500 μg/ml streptomycin and cut open using a scalpel to release the pseudocolemic fluid. The resulting fluid was centrifuged to remove any worm eggs, cells, and debris.

### Recombinant *T. muris* p43 and *T. trichiura* p47 proteins

Recombinant *T. muris* p43 (rp43) was produced in baculovirus-modified insect cells and isolated as described ^17^. Recombinant *T. trichiura* p47 (rp47) was similarly produced in insect cells (Peak Proteins (now Sygnature Discovery), Macclesfield, UK).

### Spectrofluorimetry and lipid-binding

Lipid binding by proteins was detected spectrofluorometrically in a PerkinElmer (Beaconsfield, Buckinghamshire, UK) instrument, using all-trans retinol, or the fluorescent fatty acid analogue 11- ((5-dimethylaminonaphthalene-1-sulfonyl)amino)undecanoic (DAUDA) or dansyl aminocaprylic acid (DACA) which bear the environment-sensitive dansyl fluorophore, the intrinsically fluorescent natural fatty acid cis-parinaric acid (cPnA), or the non-specific hydrophobic probe 8-anilino-1-naphthalenesulfonic acid (ANS). DAUDA and cPnA were obtained from Molecular Probes/Invitrogen (Renfrew, UK) and all other compounds were obtained from Sigma-Aldrich (Poole, Dorset, UK). The excitation wavelengths were 345 nm, 350 nm, 319 nm, and 390 nm for DAUDA and DACA, retinol, cPnA, and ANS respectively, which were at concentrations of approximately 1 μM, 1 μM, 4 μM, 4 μM, and 10 μM, respectively, in 2 ml phosphate buffered saline (PBS) pH 7.2 in a quartz cuvette. Intrinsic tryptophan fluorescence spectra were recorded for *T. muris* p43 protein in 2ml PBS with excitation wavelength of 290 nm. Emission spectra were recorded over wavelength ranges appropriate for each fluorophore to encompass peak emission in water and any shift upon entry into a binding site. Competitive displacement of fluorescent lipids was detected by a reversal of fluorescence enhancement upon addition of the ligand to a preformed complex of protein and fluorescent probe. In the fluorescence-based titration with DAUDA, its concentration checked measured using its extinction coefficient in methanol of 4400 M^-1^ cm^-1^ at 335 nm. The fluorescence spectra are uncorrected and were analyzed using Micocal/OriginLab ORIGIN software (https://www.originlab.com/). All proteins were at a stock concentration of approximately 1 mg/mL, and added to the 2 ml in amounts of 20 μl or usually less. Oleic, arachidonic acids, and other competitors were added in 10 μl quantities to the cuvettes to yield approximate concentrations in the micromolar range in the cuvette, and a series of tenfold increasing concentration increments in competitive displacement experiments.

### Protein gel electrophoresis

One-dimensional vertical sodium dodecyl sulphate polyacrylamide gel electrophoresis (SDS-PAGE) was carried out using the Invitrogen (Thermo Scientific, Paisley, UK) NuPAGE system with precast 4-12% gradient acrylamide gels, and β_2_-mercaptoethanol as reducing agent when required. Pre-stained molecular mass/relative mobility (M_r_) standard proteins were obtained from New England Biolabs, Ipswich, MA, USA (Cat. No. P7706S). Gels were stained for protein using colloidal Coomassie Blue (InstantBlue; Expedion, Harston, UK), backlit with white light and images recorded using a Nokia C32 smartphone. Colour images were converted to greyscale using IrfanView (https://www.irfanview.com/).

### Database searches and molecular structure display

Protein sequence BLAST searches were carried out through the NCBI server set to restrict to Nematoda. Sequence alignments were carried out using MultAlin (http://multalin.toulouse.inra.fr/) set for the Blossum62 substitution matrix. Amino acid content comparison plots were produced using ORIGIN from protein sequences with any cleavable secretory peptide removed as predicted by SignalP (https://services.healthtech.dtu.dk/services/SignalP-6.0/) set for Eukarya, and percent compositions calculated by ProtParam (https://web.expasy.org/protparam/). The latter were plotted against protein amino acid composition statistics from the complete UniProtKB/Swiss-Prot database release 2024_05 (https://web.expasy.org/docs/relnotes/relstat.html). PyMOL (https://pymol.org/) was used to display the *T. muris* p43 x-ray crystal structure using the RSCB PDB coordinates 6QIX.

## Results

### Fatty acid binding by *T. muris* p43

ANS was used as an initial test for binding of apolar or hydrophobic ligands, which is poorly fluorescent in water but increases its fluorescence emission dramatically upon interaction with exposed apolar sites on proteins. It has been widely used to follow folding dynamics in proteins, as a measure of the degree of unfolded protein in a preparation, or the presence of apolar pockets or binding sites. p43 bound ANS yielding a substantial increase in its emission (Figure 1A). Moreover, progressive addition of a fatty acid, oleic acid, caused a stepwise reduction in emission intensity consistent with displacement of ANS by this natural lipid. The degree of displacement was not complete, however, potentially due to the existence of apolar binding sites other than for fatty acids, or a proportion of the protein being misfolded resulting in exposure of hydrophobic internal structures.

**Figure 1.**
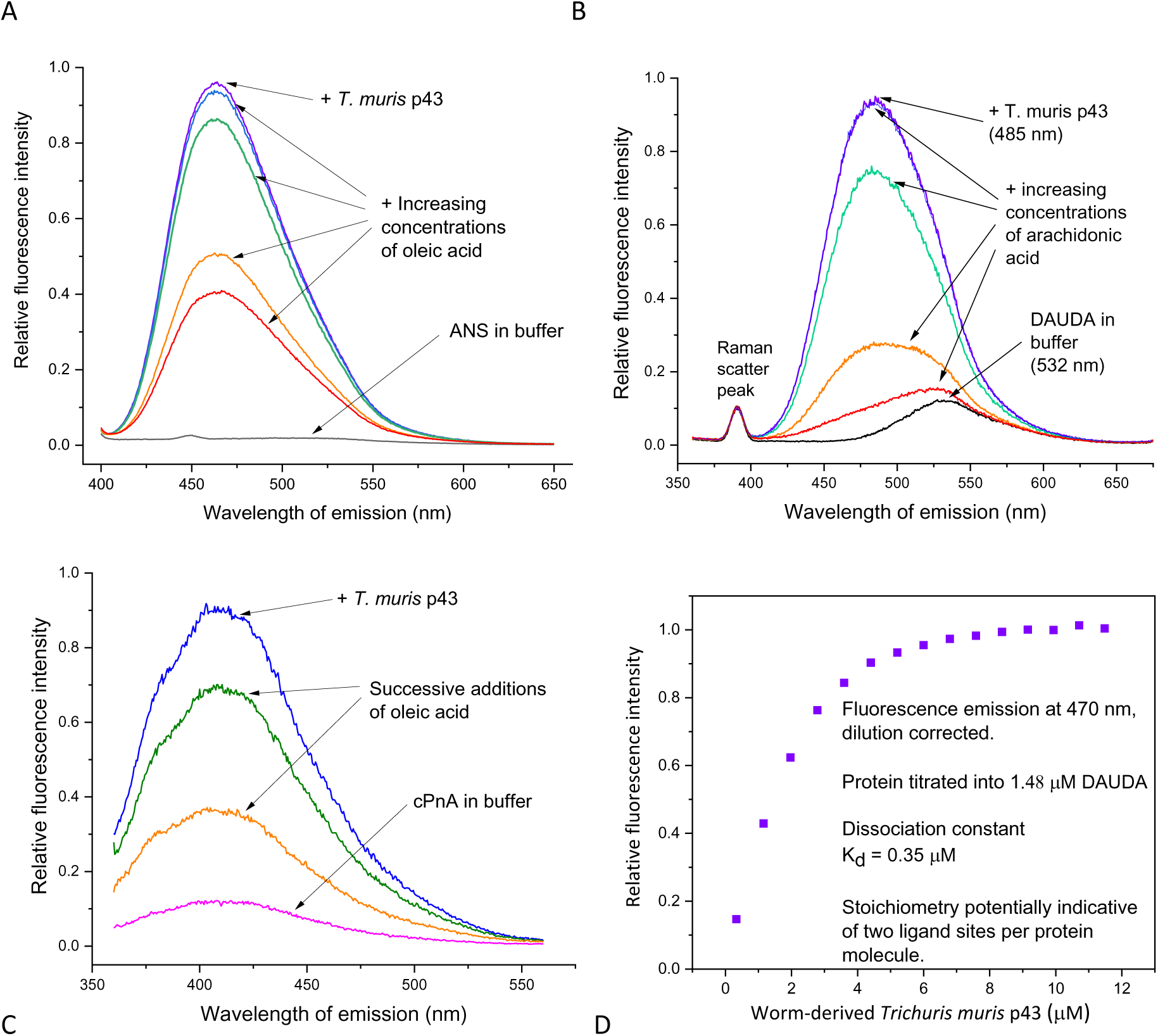
Worm-derived *Trichuris muris* p43 protein binds hydrophobic ligands. Binding was investigated by spectrofluorometry. **A,** p43 binding of the non-specific fluorescent hydrophobic probe 8-anilino-1-naphthalenesulfonic acid (ANS) to p43 and its progressive displacement by increasing concentrations of a natural fatty acid (oleic). **B,** p43 binding of a fluorophore-tagged fatty acid analogue 11-((5-dimethylaminonaphthalene-1-sulfonyl)amino)undecanoic (DAUDA) and its near complete displacement by arachidonic acid. **C,** p43 binding of a unmodified, naturally fluorescent fatty acid, cis-parinaric acid (cPnA), and its competitive displacement by oleic acid. **D,** Fluorescent titration of increasing amounts of p43 added to 1.48 μM DAUDA yielding a dissociation constant (K_d_) of 0.35 μM with stoichiometry indicative of two binding sites per protein molecule. Fluorescence intensity counts were converted to relative arbitrary units. The small sharp peaks at shorter wavelengths in B are from water Raman scatter. See Materials and Methods for excitations wavelengths for each fluorophore.

A more specific probe, DAUDA, was then used, which is a short chain fatty acid linked to a dansyl fluorophore used to examine fatty acid binding sites in proteins with more specificity than with ANS ^20,29^. The fluorophore is environmentally sensitive, poorly fluorescent in contact with water, but highly fluorescent when removed from a polar solvent in protein binding site. Upon addition of p43 to DAUDA in PBS there was a dramatic increase in fluorescence emission and, significantly, a considerable shift in peak fluorescence intensity towards the blue end of the spectrum from 532 nm to 485 nm (Figure 1B), and to a peak at 482 nm when the emission by free DAUDA is subtracted (Figure S1A). A shift of this degree is usually taken as being indicative of the probe entering a highly apolar environment such as an organic solvent ^20^, or, as here an apolar protein binding site environment or pocket. This change in DAUDA’s emission was reversible upon addition of arachidonic acid (an ω n-6 polyunsaturated fatty acid precursor to prostaglandins, prostacyclin, thromboxanes; Figure 1B), oleic acid (Figure S1B), or linolenic (another precursor to active eicosanoids; Figure S1C) acids, indicating that it was displaced from a binding site with preference for natural fatty acids. We found essentially the same observations with another fluorescent probe, dansyl aminocaprylic acid (DACA), in which the dansyl fluorophore is fixed adjacent to the carboxy end of a fatty acid rather than to the methyl/omega end as in DAUDA (Figure S1D). That DAUDA and DACA undergo similar shifts in fluorescence intensity indicated that the whole fatty acid ligand is internalised within the protein. Also, the blue shift in emission seen with DACA is even more extreme than with DAUDA (475 nm versus 485 nm, respectively), indicating that the dansyl fluorophore is taken into an even more strongly apolar environment or orientation within the binding site. The blue shift in peak emission with DAUDA and DACA is comparable to that found with other lipid-binding proteins found only in nematodes – for DAUDA, from 532 nm to 483 nm for p43, to around 472 nm with nematode polyprotein allergens (NPAs) (e.g. ABA-1, Dv-NPA-1) ^20,21^, and to about 480 nm for a nematode fatty acid and retinol-binding (FAR) proteins (e.g. Ov-FAR-1, Gp-FAR-1, Na-FAR-1) ^30–32^.

To check whether the dansyl fluorophore adduct itself contributed to DAUDA and DACA’s binding, two simpler dansyl compounds - dansylglycine and dansylamide ((5-(dimethylamino)-1-naphthalenesulfonamide) - were examined. These compounds showed only trace changes in their fluorescence spectra with p43 (Figures S1E and F) indicative of the fatty acid parts of DAUDA and DACA being the essential principles for binding to p43 rather than the dansyl group.

As a final test for specificity in fatty acid binding by p43, *cis-*parinaric acid (cPnA) was used. This is a highly unsaturated, unmodified, naturally fluorescent fatty acid with no attached fluorophore. As with DAUDA, p43 bound cPnA strongly and was displaceable with oleic acid (Figure 1C). The drop in fluorescence with oleic acid additions is much greater than due to the decay in water that cPnA undergoes with time, and the experiment was done using a concentration range of competitor that avoids confounding micelle accumulation that can occur with cPnA. The fact that p43 bound a natural fatty acid directly, and was displaced by another, is, again, strong indication of a lipid binding site or sites in p43 that are specialised for fatty acids.

### Dissociation constant and stoichiometry

A fluorescence titration experiment was carried out with increasing concentrations of p43 added to a 1.48 μM solution of DAUDA. This yielded a binding curve that reached saturation, providing an estimate of dissociation constant (K_d_) of 0.35 μM (Figure 1D). Carrier proteins that acquire then release their cargo at a destination have dissociation constants typically in this micromolar range. The calculated stoichiometry was 0.69. Under ideal conditions, a stoichiometry of 1 would indicate a 1:1 interaction, and a value of 0.5 would indicate a 1:2 protein:ligand interaction. A value of 0.69 would be consistent with two binding sites per p43 molecule in which a proportion of the protein is not binding competently, and/or that the protein contains resident ligands that are not displaced by DAUDA in the titration.

### Other hydrophobic ligands

In our previous work on nematode-specific lipid-binding proteins, retinol (vitamin A) was also shown to bind ^20,21,30–32^. Like the above probes, retinol is poorly fluorescent in water (in which it rapidly decays unless protected in, for instance, specialised proteins) and of enhanced fluorescence emission in specific, protective protein binding sites. Here, p43 bound retinol, from which it was displaceable by oleic acid (Figure 2A), indicative of a shared or overlapping binding site for retinol and fatty acids. Protein binding sites that can accommodate both fatty acids or retinol are well known in nematodes and vertebrates ^20,21,30–35^.

**Figure 2.**
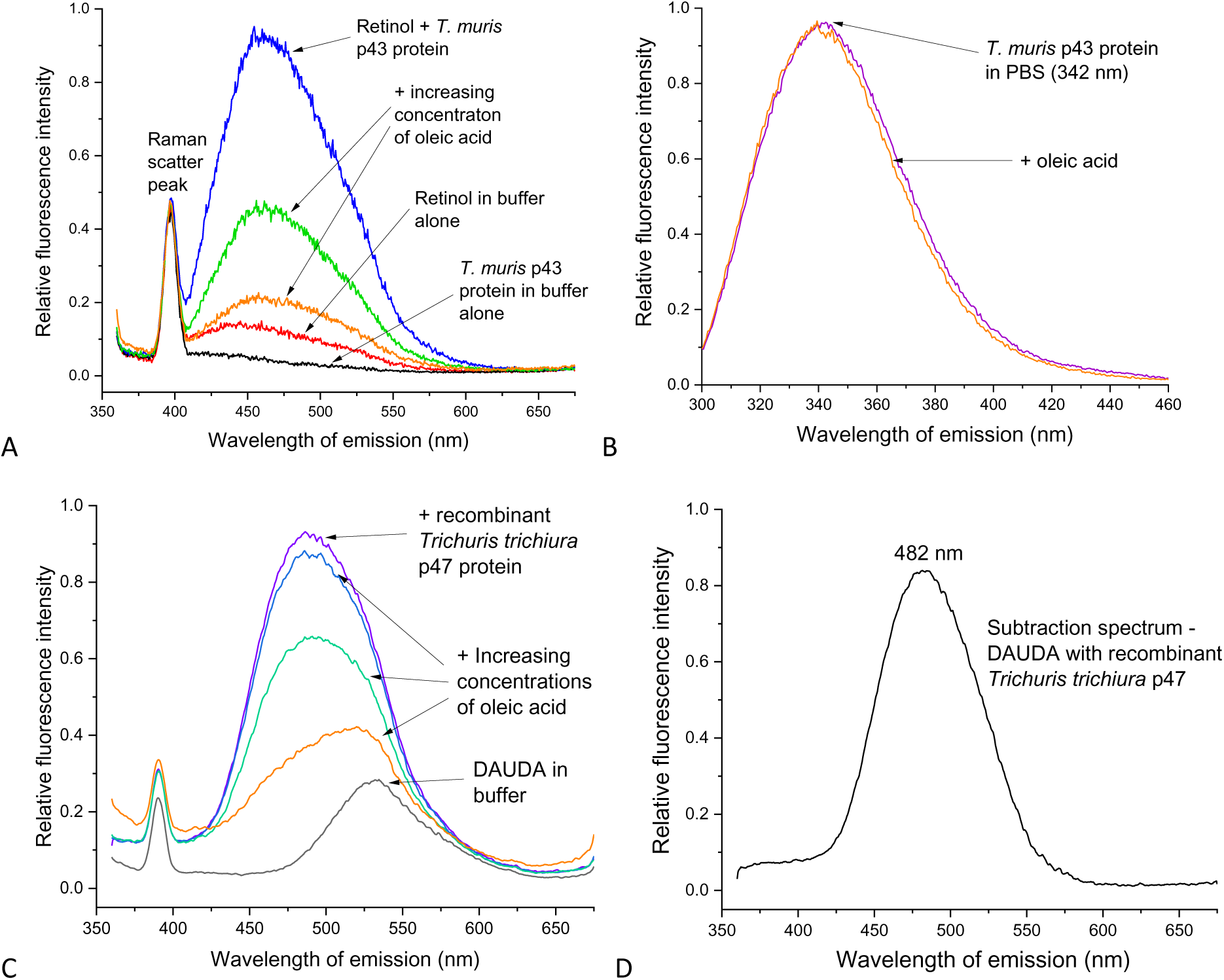
Hydrophobic ligand binding by *Trichuris muris* p43 and its orthologue from *Trichuris trichiura* p47. **A,** *T. muris* p43 binds retinol in a binding site also occupied by oleic acid. **B,** composite tryptophan intrinsic fluorescence spectrum of *T. muris* p43’s seven tryptophans with affect of addition of oleic acid. **C,** p47 of *T. trichiura,* the whipworm parasite of humans, binds fatty acids. **D,** subtraction spectrum of the spectra ((DAUDA + p47) – (DAUDA alone)) yielding peak fluorescence emission at 482 nm, essentially identical to that for *T. muris’s* p43. The small sharp peaks at shorter wavelengths in the retinol and DAUDA specra are from water Raman scatter.

Using DAUDA, other lipids were checked for binding capacity to p43, with no evidence that cholesterol displaced the probe, although this does not exclude binding in a site into which DAUDA does not partition. Similarly, the intrinsically fluorescent steroid, dehydroergosterol, did not bind p43.

To examine how the nature of the head group of modified fatty acids may influence binding, such as those that have known pharmacological/physiological proclivities, DAUDA was used in further competitive displacement experiments. Neither platelet activating factor nor L-α-lysophosphatidylcholine had more than trivial effects on fluorescence emission by a pre-formed p43:DAUDA complex (not shown). Prostaglandin E_2_ (PGE_2_) has several potent physiological effects and is secreted in abundance by *Trichuris suis* adult worms, with consequent effects on macrophage activation ^12^. However, we found very slight evidence that PGE_2_ could displace DAUDA from p43 (Figure S1G) at the pH of the buffer we used. An endocannabinoid, anandamide (arachidonylethanolamide), that has several pharmacological effects, including in the gut ^36–40^, showed evidence of displacing DAUDA but to only a limited degree (Figure S1H). Anandamide comprises arachidonic acid with a relatively small change in the size of its head group – from a carboxylate to an ethanolamine, ethanolamide head group – yet arachidonic acid itself displaces DAUDA from p43 highly effectively (Figure 1B). So, despite the fact that several (polyunsaturated) fatty acids can bind p43, there is a degree of specificity defined by the headgroup. Again, both PGE_2_ and anandamide may, however, bind to a subset of binding sites in p43 into which DAUDA may not enter.

### Intrinsic tryptophan fluorescence

Proteins contain a few intrinsically fluorescent amino acids, the most useful of which is tryptophan (Trp), the emission spectrum of which provides information on the environment of its indole side chain ^41^. This, a Trp that is fully exposed to solvent water in a fully unfolded protein will exhibit a peak fluorescence emission at approximately 356 nm, and 318 nm in one that is fully buried. The peak Trp emission of p43 occurred at 342 nm, which will be a composite of the emissions of its seven Trps in their diverse environments (Figure 2B). When oleic acid was added to p43 in buffer there was a slight blue shift that may occur upon a ligand entering a binding site where a Trp were present, or that ligand binding causes an alteration in the conformation of the protein that impinges on the local environment of one or more distal tryptophan sidechains.

### Lipid binding by the p43 orthologue of the human parasite, *Trichuris trichiura*

Given the close phylogenetic relationship between *T. muris* and the similar parasite of humans, *T. trichiura,* we examined lipid binding using a recombinant version of the latter’s p43 orthologue, p47. The amino acid sequences of the proteins absent of their cleavable N-terminal signal sequences (cleavage site between position 17 and 18 (as predicted by Signalp 6.0) and terminating at the same place, are 84% identical. The *T. trichiura* p47 bound DAUDA in an essentially identical manner to *T. muris* p43 in terms of fluorescence enhancement, blue shift in peak fluorescence emission, and displacement by oleic acid (Figure 2C). A subtraction spectrum to isolate emission by DAUDA when within the protein yielded a peak of fluorescence emission at 482 nm (Figure 2D), which is essentially indistinguishable from *T. muris* p43 (Figure S1A).

### p43 dominance in *T. muris* pseudocoelomic fluid

In multicellular animals, abundant extracellular lipid-binding proteins are often associated with resource distribution systems such as the blood of vertebrates (e.g. serum albumin, the components of lipoprotein complexes such as LDL, HDL, chylomicrons), and the haemocoel of arthropods (e.g. lipophorins of insects. Very little is known about the PCF of parasitic nematodes, except in the few large species from which it can be easily extracted such as *Ascaris suum*, the large roundworms of pigs, *Toxocara* canis, the roundworm of dogs, and *Dioctophyme renale,* the giant kidney worm (many host species) ^20,22,26^. Of these, the latter is in the same sub-clade of the initially genomically-designated Clade I of nematodes (the Dorylaimia) as *Trichuris* ^27,28,42^. The PCF of *D. renale* has two proteins that are abundant, P17 (a bright red, haem-bearing protein and presumed globin) and P44 (a fatty acid-binding orthologue of *T. muris* p43). SDS-PAGE comparison of the PCF of *T. muris* and *D. renale* showed a similar dominance of p43 and P44 in the respective parasites, but with p43 seemingly being considerably the more abundant relative to a ∼17 kDa protein (the possible globin orthologue of P17 of *D. renale*) than in *D. renale* PCF (Figure 3). p43 migrates more slowly in the electrophoresis gels than does P44, presumably because of the absence of a histidine-rich C-terminal sequence in the latter and possible differences in glycosylation. Both p43 and P44 migrated more slowly when reduced, which is also seen in mammalian serum albumin, and is presumably due, in the case of both p43 and P44, to the cleavage of the unusually high number of internal disulphide bonds causing unravelling of the molecule and thereby impeding its migration through the gel.

**Figure 3.**
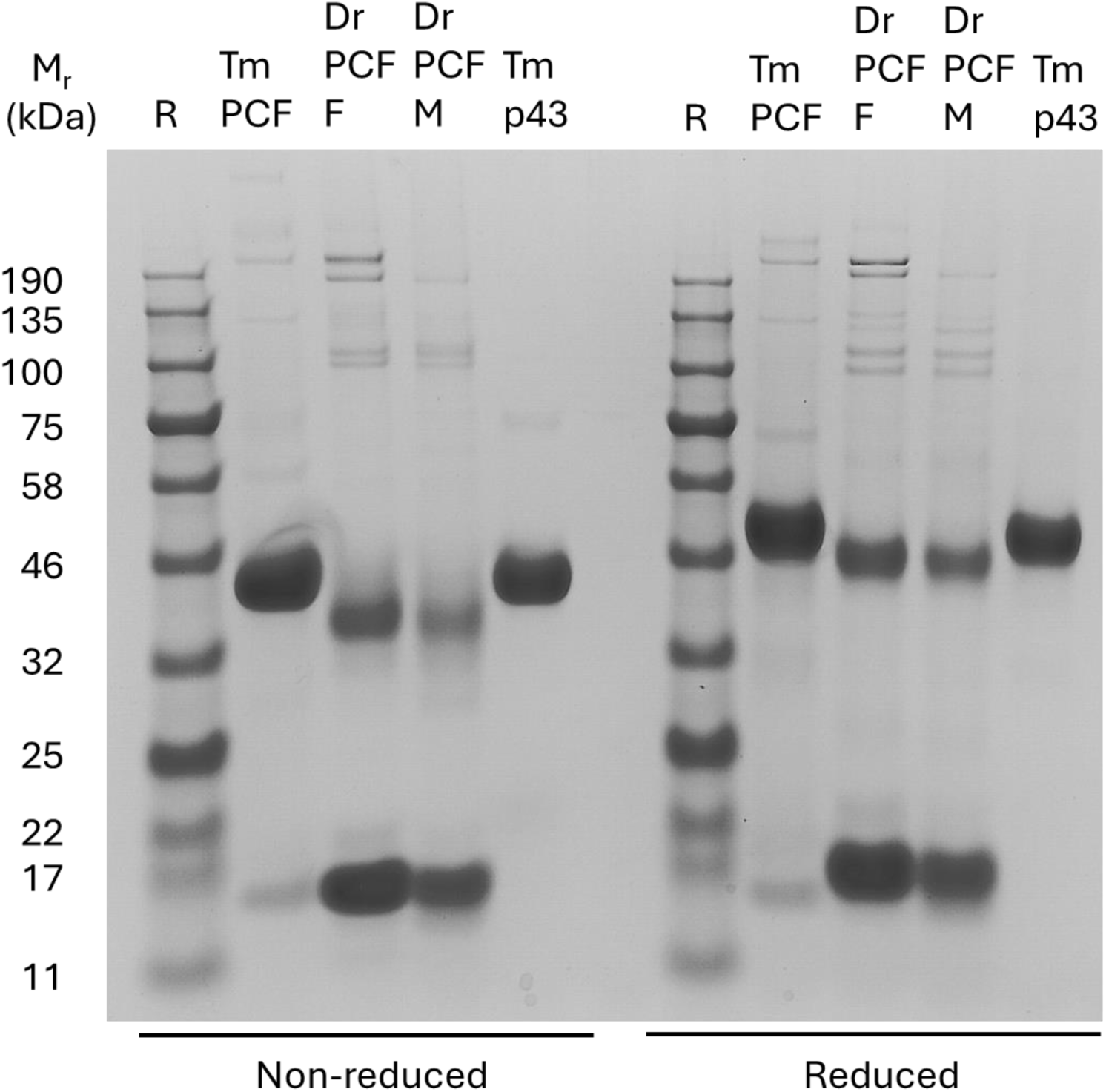
p43 of *Trichuris muris* is the dominant protein of the worm’s pseudocoelomic fluid (PCF), as is its orthologue in a related parasite. The two species are in a subbranch of Clade I nematodes, the Dorylaimia ^28^. Protein electrophoretic comparison of the PCF of *T. muris* and the giant kidney worm, *Dioctophyme renale*, along with purified, worm-derived *T. muris* p43. The reduction in mobility of p43 and p47 under reducing conditions is typical of proteins with extensive internal cysteine-cysteine cross-links that are lost upon reduction, likely resulting is unravelling of the protein that impedes migration through a gel. 4-12% gradient SDS-PAGE. Coomassie blue stained. M_r_, relative mobility of standard reference proteins (R) expressed in kiloDaltons (kDa). PCF, pseudocoelomic fluid; F, female; M, male; Tm, *Trichuris muris*; Dr, *Dioctophyme renale*.

### Potential ligand-binding cavities within the structure of p43 and their diversity

The only published structure of a dorylipophorin is that of *T. muris* p43. The protein’s amino acid sequence clearly indicates an internal duplication (see alignment in Figure S2; and such that it comprises two highly integrated sub-domains that are further linked by a disulphide bond, at the junction of which is predicted from molecular simulations to be the IL-13-binding site ^17^. The protein’s structure is novel, and does not resemble the structures of the two other nematode-unique classes of lipid-binding proteins found in Clades III, IV, and V (the Chromadoria), namely the nematode polyprotein allergens (NPAs)^43^ and the fatty acids and retinol-binding proteins (FARs) ^31^. p43 possesses unusual features such as a remarkably high proportion of Cys, Lys, Pro, and His residues, even excluding the extensive His-rich tail (Figure S3). The network of internal disulphide bonds potentially stabilises the structural integrity of a protein that comprises an extensive set of internal cavities that vary in the charges of their walls (Figure 4). Notably, the size, shape, and surface charge of the equivalent cavities can be seen to differ between the two sub-domains. If the x-ray diffraction structure of p43 is shown absent of the protein structure itself, many of the cavities can be seen to be occupied by components of the crystallisation buffer, indicative of accessibility to the outside of the protein’s structure (Figure S). Some of the cavities have features reminiscent of lipid-binding cavities in other proteins such as serum albumin of vertebrates ^44,45^, and the FAR and NPA proteins of nematodes ^31,43,46^, namely an elongated tunnel with predominately apolar walls and sometimes a charged end that may act to tether amphipathic compounds such as fatty acids and retinol. Other cavities in p43, however, have quite different surface charges so may bind different ligand types with quite different charge characteristics.

**Figure 4.**
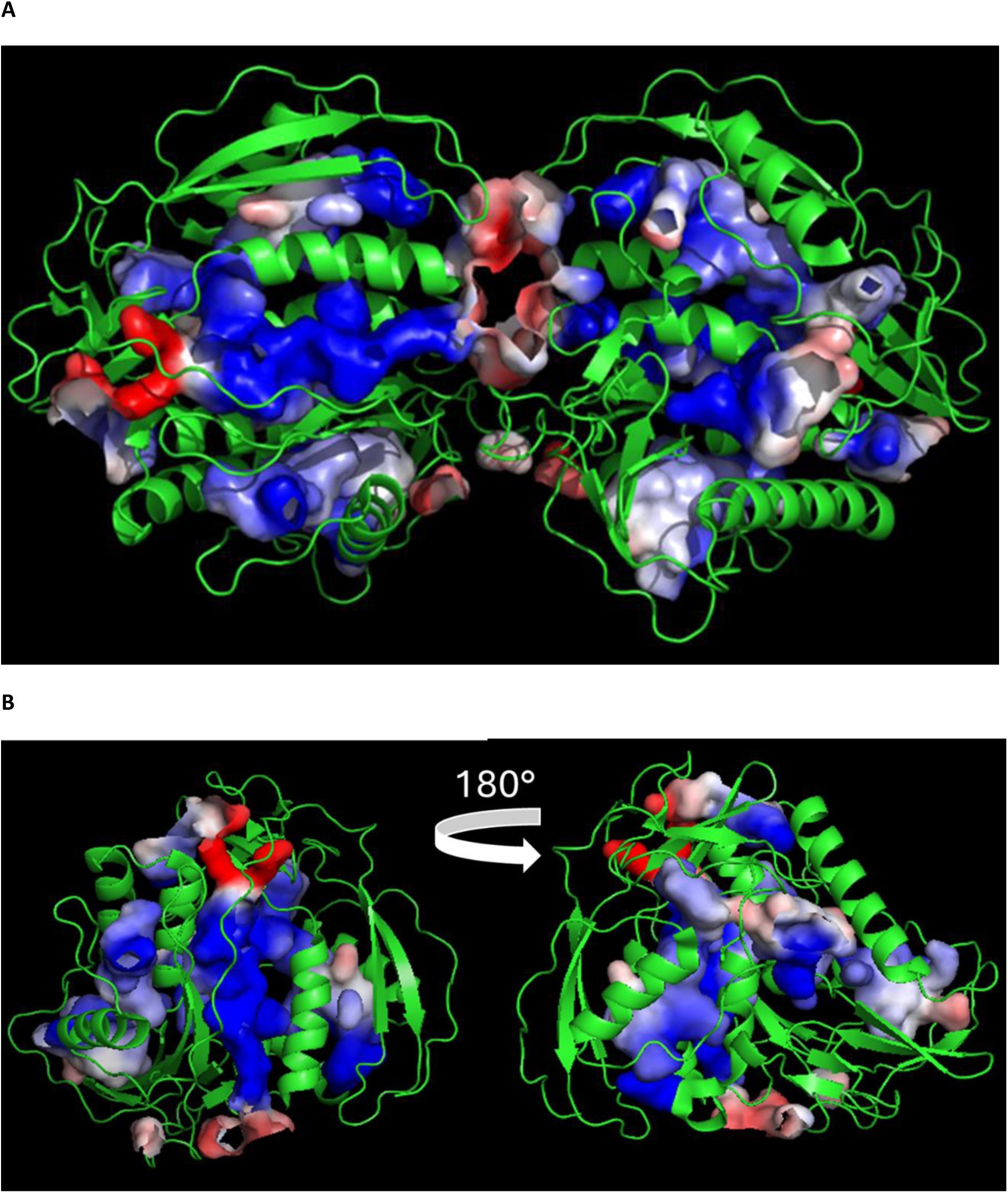
*Trichuris muris* p43 protein possesses extensive surface-connected internal cavities. Ribbon representation of the X-ray crystal structure of p43 showing a mix of alpha helices (coils) and extended/β structures (arrows) along with unstructured chains. **A,** Two molecules as in crystal structure asymmetric unit (PDB 6QIX); cartoon ribbon for chains; cavity surfaces coloured for electrostatic potential of their walls, culled to show only internal cavities; charges indicated - red for negative, blue positive, white neutral. Cavities exhibit a complex mix of internal surface charges and shapes, and the cavities of the duplicated/repeat domains appear to have diverged. **B,** Rotation of one molecule, again showing the diversity of cavities and the lack of repetition of types. Molecular display using PyMOL.

### Amino acid sequence diversity of Clade 1 p43 orthologues

There are currently complete amino acid sequences for DLPs from three Clade I nematode genera, namely *Trichuris, Trichinella,* and *Dioctophyme,* along with several incomplete sequences from *Soboliphyme baturini.* An alignment of the three complete sequences reveals distinct regions of similarity, notably around the highly conserved cysteines, interspersed with regions of divergence (Figure S5). If the regions of conservation are mapped onto p43’s structure, they tend to associate with extended/β structure and external loops, whereas the helical regions tend to be less conserved (Figure S6). This may indicate that surface features of DLPs are involved in interactions with entities such as other proteins, membranes, or membrane receptors that may be common to species within Clade I nematodes. That the DLPs of *Trichuris* and *Trichinella* possess long histidine-rich tails, but that of *Dioctophyme renale* does not, hints at some differences in function of DLPs that may include, in the case of parasites, animal or plant, how they interact with their hosts.

## Discussion

We have shown that the major secreted protein of the whipworm *T. muris*, the highly immunomodulatory p43 protein, is a lipid-binding protein. It can bind polyunsaturated fatty acids such as arachidonic and linoleic acids that can themselves be pharmacologically active or be the precursors of biologically active eicosanoid lipids that include prostaglandins, leukotrienes, thromboxanes, and several other classes of signalling lipid ^47–51^. p43’s lipid binding function may therefore add to controlling immune and inflammatory processes at the infection site and beyond either by delivering, or sequestering, specific lipids in order to compromise and modify the host’s own cellular signalling pathways. Notably, we also found that a recombinant form of the p43 orthologue (p47) in the human-parasitic whipworm *T. trichiura* has similar binding properties. p43 also binds retinol, which is a precursor to a wide range of retinoids that are important signals in numerous immune, cellular activation, differentiation, and repair responses ^51–56^.

As can be seen from Figure 4, *T. muris* p43 has numerous internal cavities that communicate with its surface. Some that do not so communicate visibly in the x-ray structure may do so in solution when the protein’s structure may be dynamic. The shapes of some these cavities, and the charge distributions of their walls, could be consistent with occupancy with hydrophobic ligands, or of binding compounds with different charge characteristics to lipids. A superficial visual comparison of the cavities and pockets of p43’s two repeat domains indicates that the domains may have diverged in their binding functions (Figure 4). The same can be said of vertebrate serum albumin, which is also constructed of repeat domains (three) with several lipid-binding pockets, and, like p43, is held together by an unusually large number of disulphide bridges for a protein of its size ^44,45^.

The first p43-like protein was defined from *Trichinela spiralis* and termed poly-cysteine and histidine-tailed protein (PCHTP) because of its unusual richness in these amino acids, as is p43 (Figure S3A) ^17,25^. The His-richness derives from both their main sequence and an extended His-rich carboxy terminal tail. The proposed collective systematic name for these orthologues, “dorylipophorins” (DLP), is a portmanteau reflecting their confinement to the Dorylaimia (Clade I) nematodes, and their now-known biochemical activity. Histidine-richness is not always excessive in these proteins – the DLP of the giant kidney worm, *D. renale*, for instance, does not exhibit a His-rich carboxy terminal tail, or such a His-rich main sequence as does p43 (Figure S3A & B). Differences in His-richness of these proteins could be relevant to the differing biologies and parasitologies of this group of nematodes. Histidines can be associated with proteins that bind divalent cations (e.g. zinc, nickel, copper, iron) or haem, as in histidine-rich glycoprotein ^57,58^, an abundant protein in human plasma, haemoglobin ^59^, and the histidine-rich, haemozoin-binding protein of *Plasmodium* malaria parasites ^60,61^. Notably, for p43, zinc is contingent to the protein’s interactions with glycosaminoglycans such as heparin sulphate ^17^, which may be an aspect of its biological activity that may apply to other parasites. Curiously, His-rich regions in major pseudocoelomic lipid-binding proteins of nematodes distantly related to *Trichuris* are apparent (e.g. *Dictyocaulus viviparus’s* Dv-NPA-1 ^24^; and *Caenorhabditis elegans’s* NPA (gene CELE_VC5.3)), so metal- (and possibly heme-) binding may be a common feature of some internal transport proteins in nematodes.

The extraordinary abundance of cysteines in p43 and other DLPs is also of particular note ^17,25^; Figure S3) and may relate to the need to hold together a protein structure with such extensive intramolecular spaces, particularly if it were to be a dynamic structure. The elevated content of prolines in DLPs (Figure S3), p43 in particular, may also relate to this. The possibility also exists that binding of different lipids could modulate the structure of p43 to materially alter its biological activity, of which there are examples such as the surface spike protein of the SARS-CoV-2 virus ^62,63^.

The dominance of p43 in the PCF of *T. muris,* along with that of its orthologue in *D. renale,* argues that the original function of DLPs was as internal bulk lipid transporters in the body fluid of this clade of nematodes, akin to serum albumin and the lipoproteins of vertebrates, and the lipophorins of insect haemolymph. DLPs do not appear to exist in Clades III, IV, and V nematodes (the Chromadorea of the original DNA-based classification ^64^, subsequently refined ^27,28,42^), their functions possibly replaced by other internally abundant lipid-binding protein classes found only in those nematodes, specifically, the NPA and FAR proteins, that are in turn absent in the Dorylaimia. There is currently no information on pseudocoelomic lipid binding proteins in the remaining branch of the nematodes, Clade II, the Enoplea, and we have so far found no signs of DLP-, NPA-, or FAR-like protein sequences in searching the albeit limited genomic information on this clade. Intriguingly, therefore, two, and perhaps three, branches of the Phylum Nematoda evolved structurally distinctive proteins, likely with no common ancestor protein, to perform a critical function in multicellular animals, that of distributing water-insoluble, and sometimes chemically unstable, lipids in bulk internally. In some of the Dorylaimia parasitic forms at least, here exemplified by *T. muris*, the DLPs may have been further adapted for external roles such as interactions with a parasite’s host’s tissues. Intriguingly, in addition to all larval and adult stages, p43 is also found in embryonated eggs of *T. muris* ^65^, so it may play a part in lipid storage and mobilisation for the developing embryos, and maintenance of eggshell integrity such as is postulated for a fatty acid binding protein found in the perivitelline fluid of *Ascaris* ^66,67^. It is of course likely that dorylipophorins could play a similar role in free-living members of the clade.

It remains to be seen whether the lipid-binding activity of p43 has any bearing on its immunomodulatory activity. However, one eicosanoid lipid, PGE_2_, has been demonstrated to be released in large quantities by adult worms of another *Trichuris* species, *T. suis* and the worm-derived PGE_2_ can suppress proinflammatory properties in human dendritic cells ^12^. PGE_2_ is similarly secreted by *T. muris* (Bancroft and Grencis, unpublished). We did not, however, find good evidence that p43 binds PGE_2_ in our particular fluorescence-based competition assays, so it may not need a carrier protein for delivery to host tissue. The water-solubility and chemical stability of PGE_2_ is pH-dependent ^68^, however, so it may partition into a carrier protein at some stage in its synthesis and export. Meanwhile, there is increasing evidence that lipids are important in affecting immune activities in the gut, cases of which are relevant to anti-helminth immunity ^36,51,69–72^.

There is potentially a difference between compounds that p43 can bind experimentally and that with which it is loaded by the parasite internally or acquires externally *in vivo*. In a preliminary lipidomics analysis, we extracted lipids from the worm-secreted p43 used in this study and found the eicosanoids eicosapentaenoic acid (EPA), docosahexaenoic acid (DHA), arachidonic acid (AA), 5,6 epoxyeicosatrienoic acid (5,6-EET), and 15-hydroxyeicosatetraenoic acid (15-HETE) (M.W. Kennedy, A,J, Bancroft, C. Regnault, P. Whitfield, R.K. Grencis, unpublished). The latter two compounds require enzymatic modification of a polyunsaturated fatty acid precursor to confer biological activity and perform a wide range of signalling functions ^73–76^. These preliminary findings suggest that either the parasite has eicosanoid-modifying enzymes similar to its host (*T. suis* appears to; ^12^), or that these lipids are harvested from the host and then released in p43 to modify the immunological and inflammatory environment of the infection site. We also found palmitic acid, a saturated fatty acid involved widely in energy metabolism and membrane construction – this is unlikely to be involved in immunomodulation but is consistent with a function for p43 in internal lipid distribution of an otherwise insoluble metabolic component. These binding activities of *T. muris* p43 are consistent with a dual role in distribution of lipids internally for the worm’s own physiological needs, and externally for environmental modification. Indeed, it remains conceivable that these nematodes may use eicosanoid lipids for their own cell-cell signalling purposes.

*T. muris* p43 (and presumably also p47 of *T. trichiura*) has joined a widening array of protein types that are secreted by parasitic nematodes to control and alter their environments. A curiosity is that these include lipid-binding proteins with similar binding propensities – p43 for *T. muris* at the very least for Dorylaimia species, and the FAR proteins in Chromadoria - FAR proteins have been found to be influential in both plant and animal parasitism ^77^. P43 orthologues may similarly be important in plant, vertebrate, and arthropod (for example the Mermithids of insects) parasitism by other members of the Dorylaimia. For the moment, our findings of p43’s lipid-binding activities, and a future better understanding of its binding sites for lipids and perhaps other ligands, opens the way to test mutated versions of the protein in immunomodulation and control of the infection site, by, for example, investigating the effects of modified p43 on immune cells *in vitro*, or the success or otherwise of p43-genetically modified parasites *in vivo*.

### Supplementary/supporting information is available at the end of this document Acknowledgements

We are most grateful to Gisela Franchini of La Plata University, Argentina, who kindly provided a sample of pseudocoelomic fluid from *Dioctophyme renale,* and to Alan Cooper for advice on data interpretation.

## Author contributions

Conceptualization, M.W.K., R.K.G., A.J.B.; Experimentation, data curation, formal analysis, M.W.K.; Funding acquisition, R.K.G., M.W.K.; Methodology, M.W.K., R.K.G., A.J.B.; Project administration, M.W.K., R.K.G., A.J.B.; Supervision, M.W.K., R.K.G., A.J.B.; Visualization, M.W.K.; Writing original draft, M.W.K.; Writing review & editing, M.W.K., R.K.G., A.J.B..

## Data availability

The original data presented in the study are included in the article or Supplementary Materials. Further inquiries can be directed to the corresponding authors.

## Supplementary/supporting information

**Figure S1.**
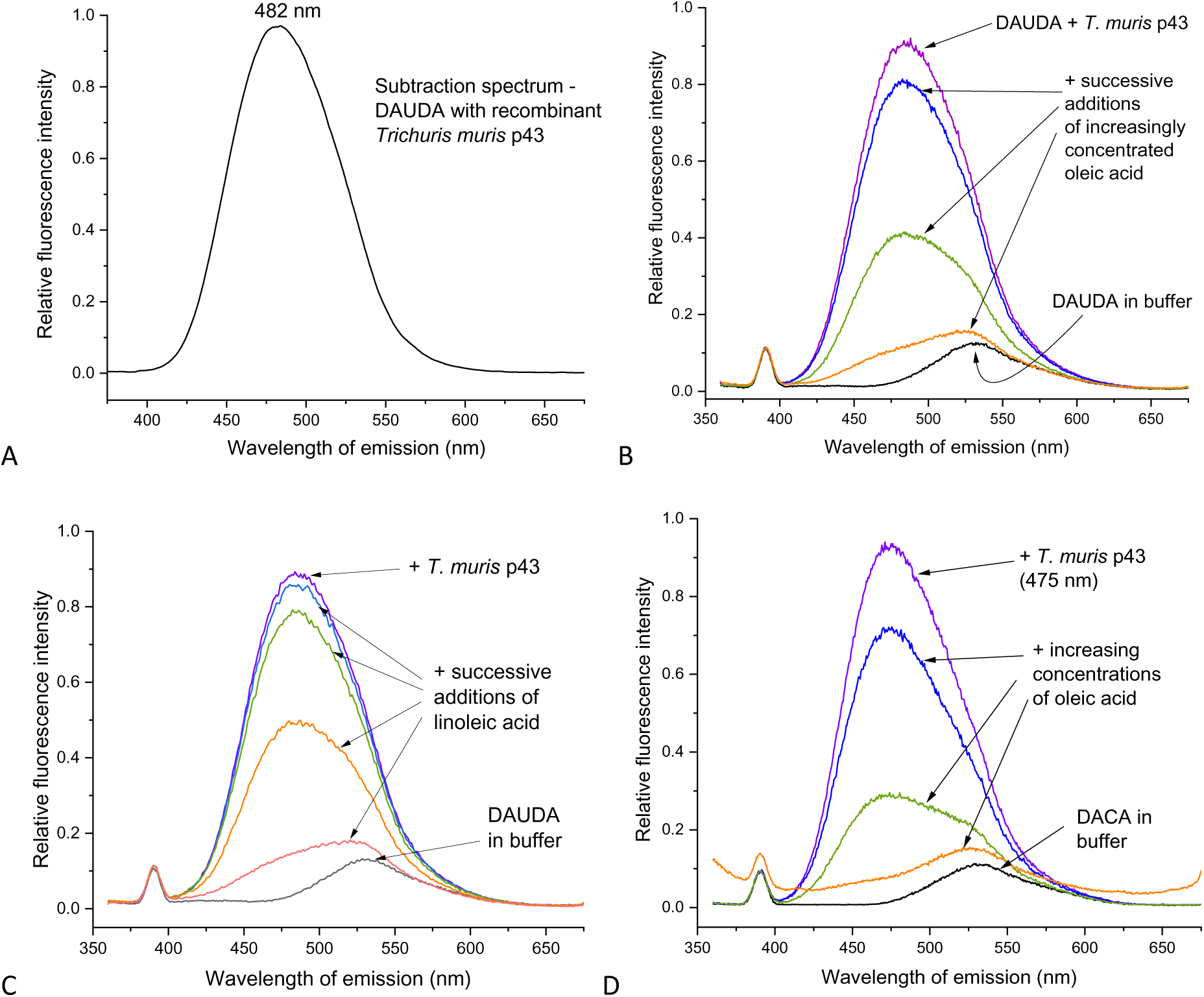
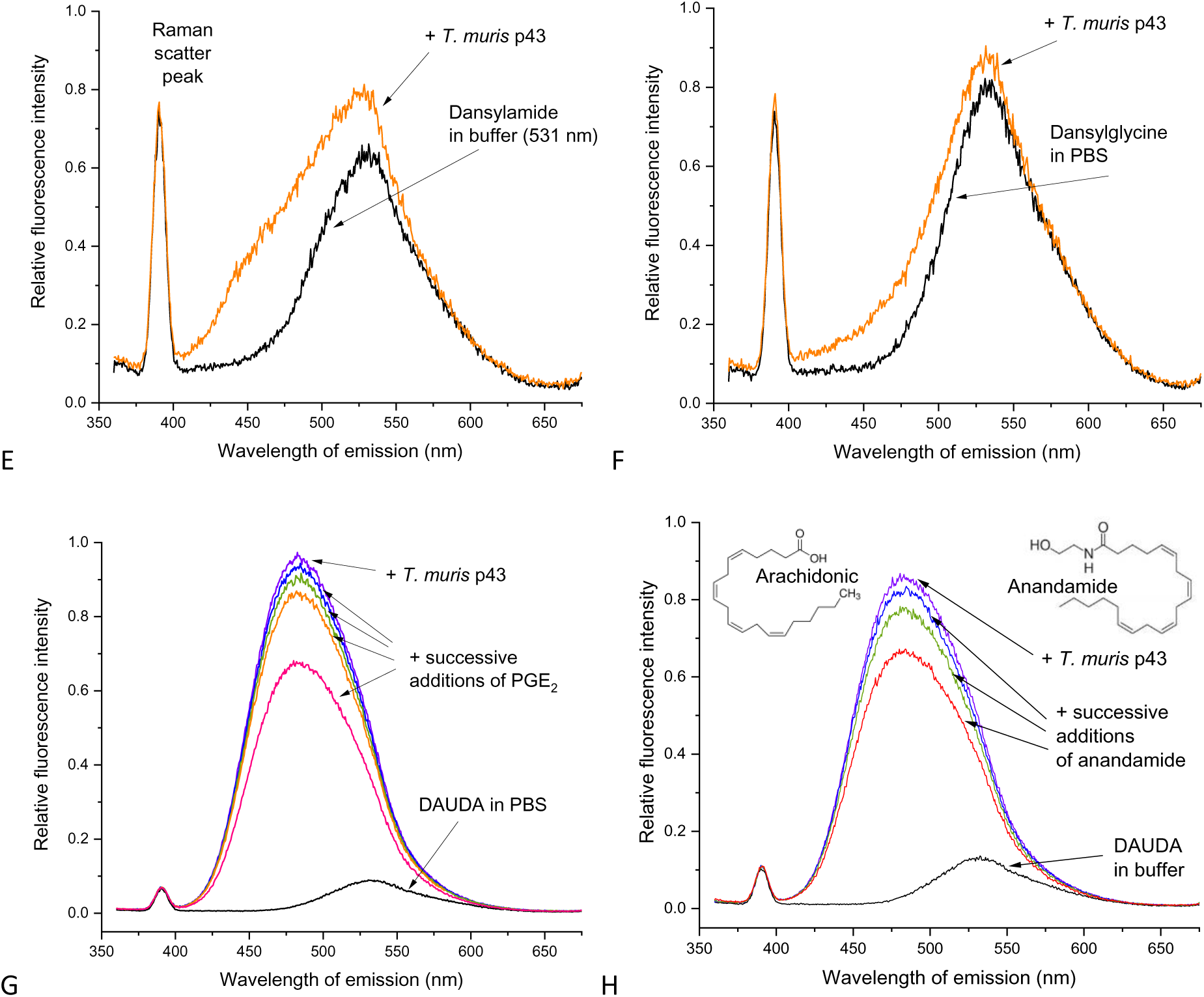
Hydrophobic ligand binding to the immunomodulatory p43 protein of *T. muris*, and likely irrelevance of an attached dansyl fluorophore. **A,** subtraction spectrum ((DAUDA + *Trichuris muris* p43) – (DAUDA alone in PBS)) of DAUDA binding to p43 showing the strong blue shift in peak fluorescence emission of the dansyl fluorophore (from 532 nm to 482 nm) when in the apolar environment of the protein’s binding site. This shift is identical to that for *T. trichiura’s* p47 protein (see main text). **B,** near complete competitive displacement of DAUDA from p43 by oleic acid. **C,** as for **B** but with linoleic acid (like arachidonic acid, a polyunsaturated fatty acid precursor to many biologically active eicosanoid lipids). **D,** binding of dansyl aminocaprylic acid (DACA) to p43 and displacement by oleic acid. DACA bears the same dansyl fluorophore group as does DAUDA, but it is attached by the carboxylate headgroup of the fatty acid instead of at the methyl end of the hydrocarbon tail. This is consistent with the entire fatty acids with their respective attached dansyl group being taken into the binding site. **E, F,** lack of binding of two small dansylated compounds to p43, indicating that the dansyl group itself does not contribute to binding by DAUDA or DACA. **G, H,** meagre displacement of DAUDA bound to p43 by either prostaglandin E_2_ or anandamide. Anandamide comprises arachidonic acid (which does displace DAUDA strongly) with an ethanolamide head group. So, a relatively minor change in headgroup can interfere with lipid binding.

**Figure S2.**
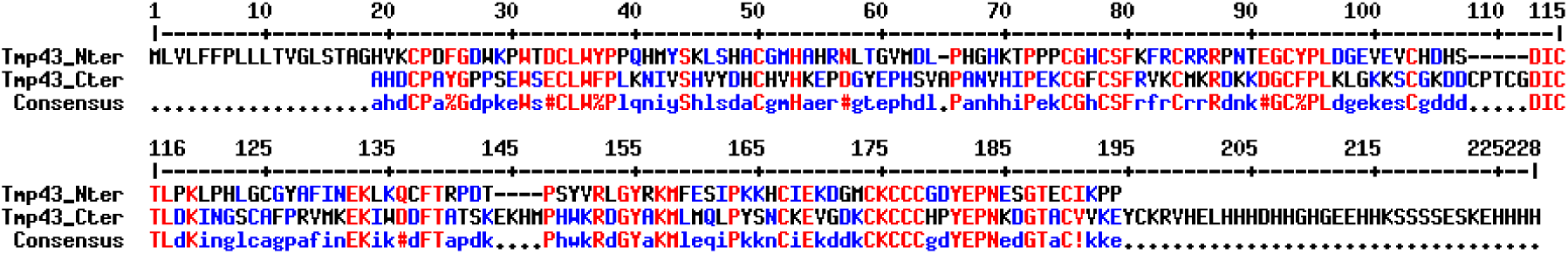
*Trichuris muris* p43 internal duplication. Alignment of N- and COOH-terminal halves of *T. muris* p43. SignalP 6.0 predicts cleavage of secretory signal sequence between position 17 and 18 of the nascent protein. Alignment made with MultAlin, as per Figure S2.

**Figure S3.**
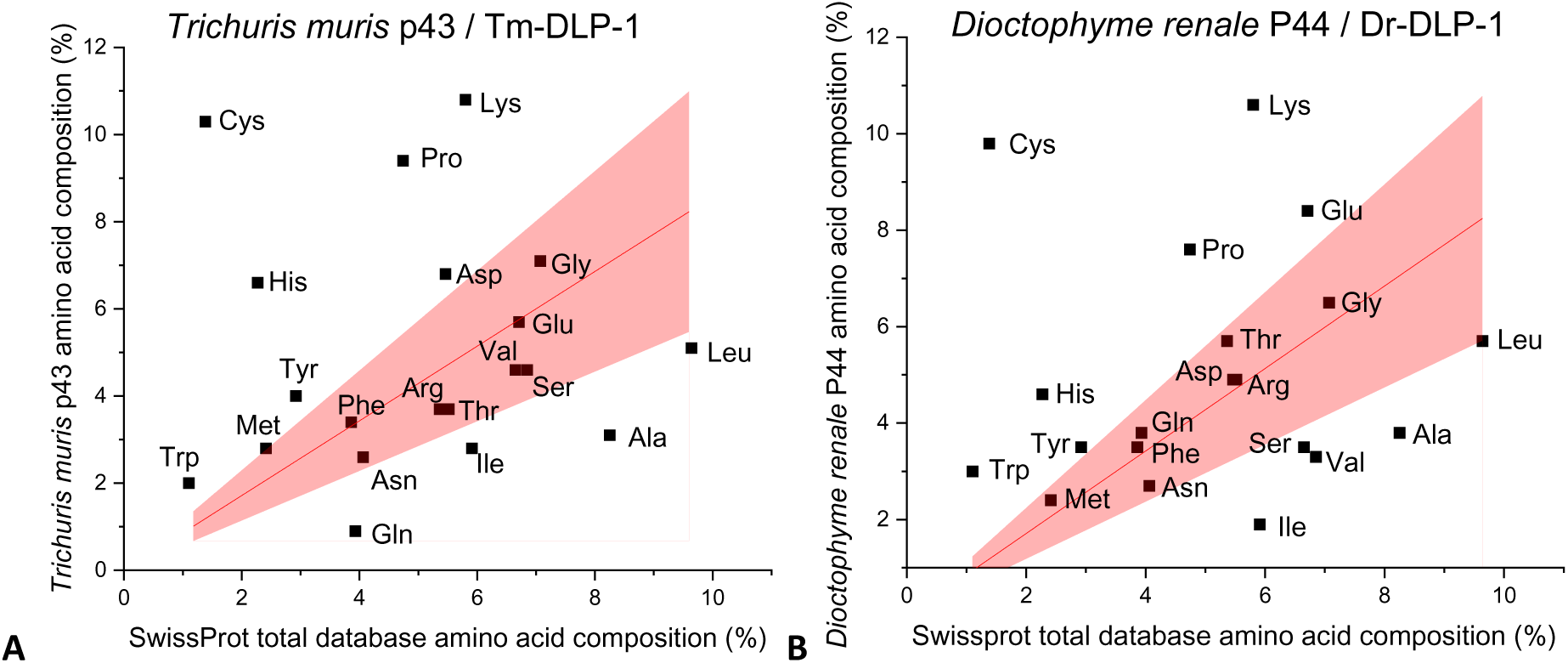
Comparison of the amino acid compositions of *Trichuris muris* p43 and the orthologue from *Dioctophyme renale* (Dr-DLP-1) against the global average composition of all entries in the SwissProt protein database. (https://web.expasy.org/docs/relnotes/relstat.html). **A.** *T. muris* p43, cleavable secretory signal sequence as predicted by SignalP removed, so count began from position 18 at amino acids GHV in the sequence; count truncated just before the histidine-rich tail at position HEL so as properly to compare the main sequences of the two proteins. **B,** The same analysis for Dr-DLP-1/P44 of *D. renale,* which does not have a long histidine-rich COOH-terminal tail. This illustrates the near-identical enrichment with cysteines in both proteins and above average histidine richness typical of this family of proteins even excluding any histidine-rich tails (such as in in *Trichuris* and *Trichinella* species *spp.* and *Trichinella spp.*) although the level of His-richness of Dr-DLP-1 is less apparent. Linear curve fits; intercept set for zero; 95% confidence bands coloured. The unusual proline richness in both proteins may relate to maintaining crucial structural elements, though again, p43 shows this more than does Dr-DLP-1. *T, muris* p43 sequence from Genbank NCBI 6QIX_A; *D. renale* Dr-DLP-1 sequence from GenBank NCBI QTE33903.2

**Figure S4.**
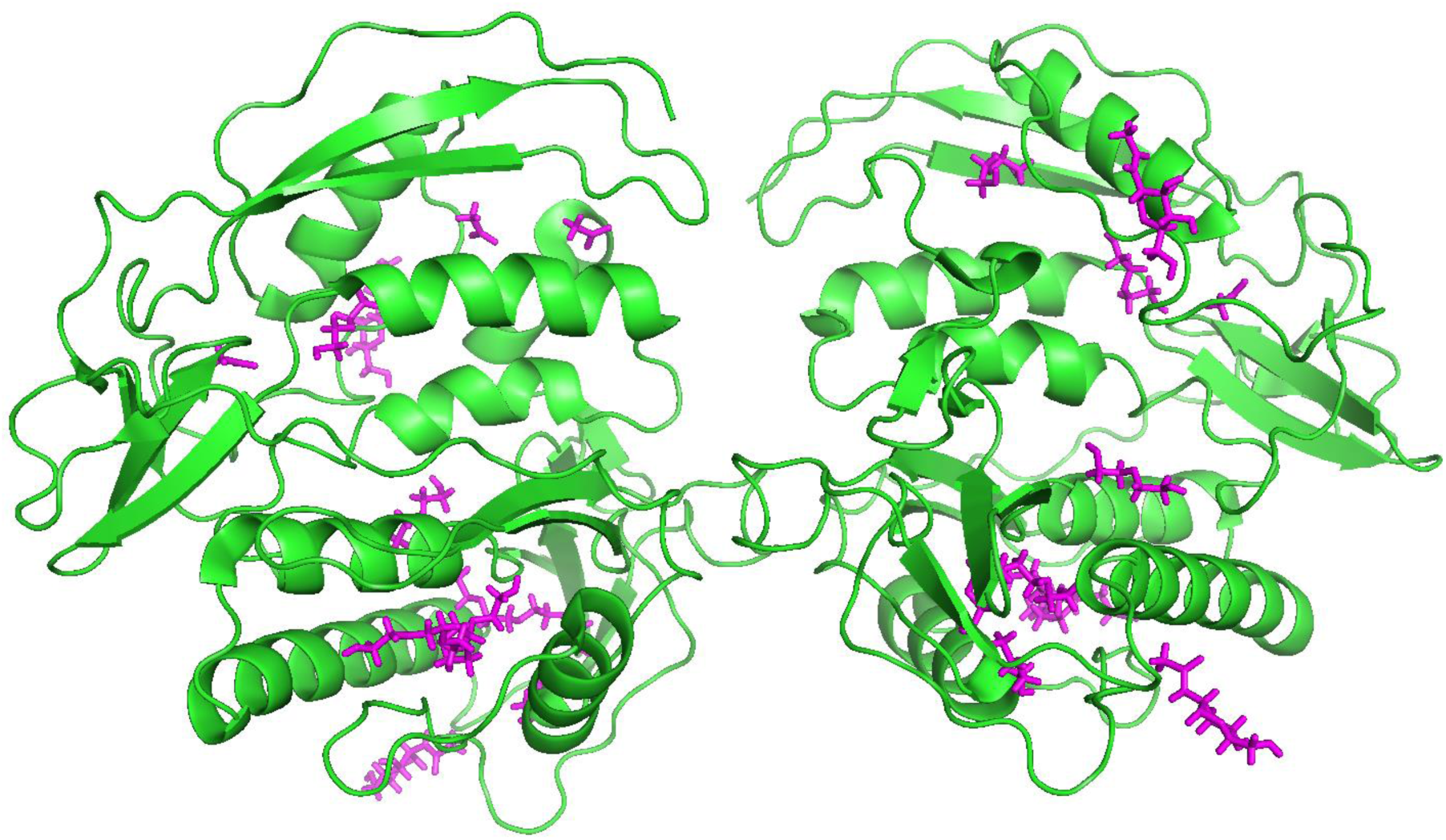
Surface accessibility of internal cavities of p43. X-ray crystal structure of p43 (PDB 6QIX_A) showing the two molecules of the asymmetric unit with the components of the crystallisation buffer *in situ* (magenta sticks), some of which are on the surface of the protein, but many are internal.

**Figure S5.**
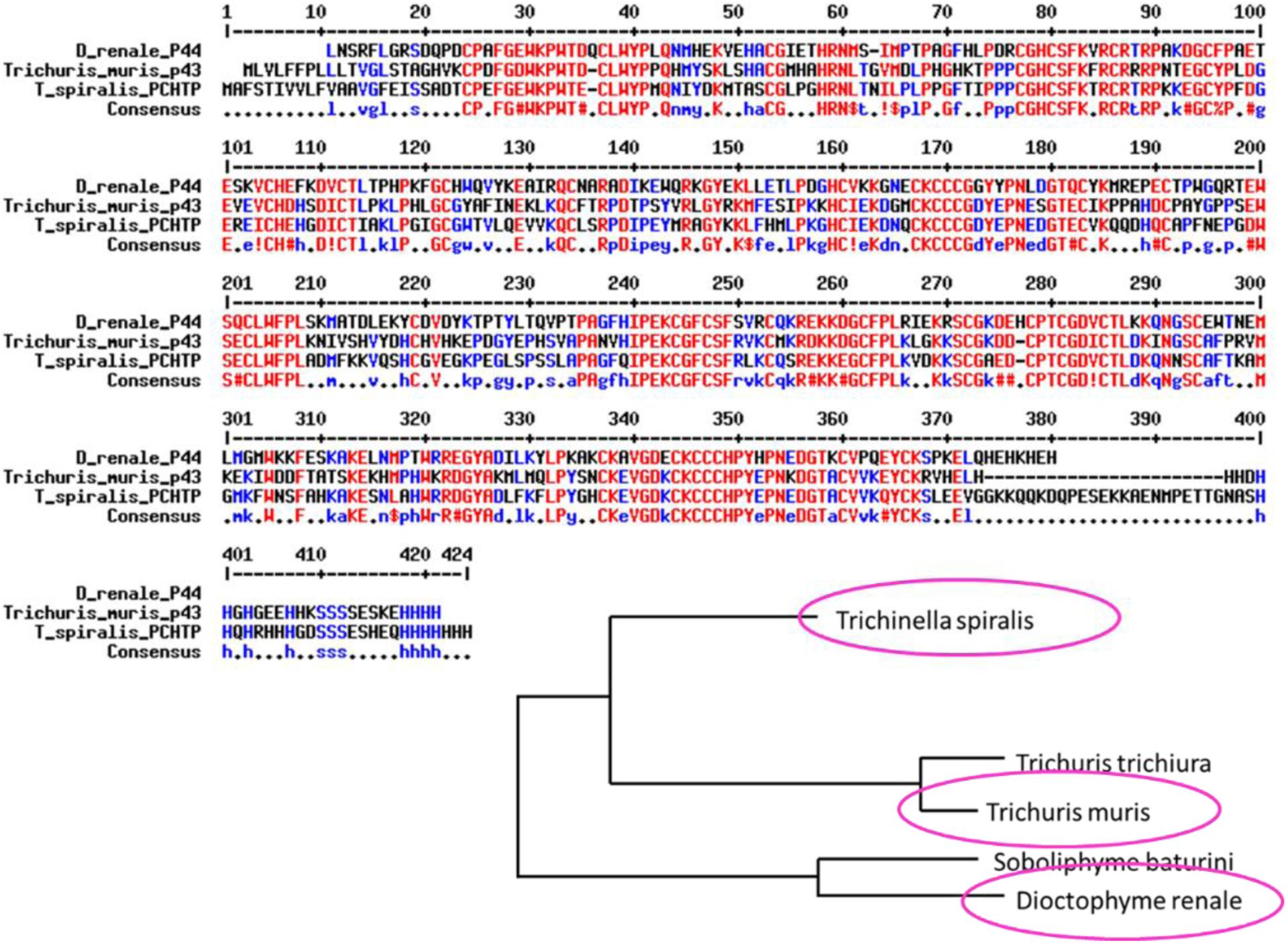
Alignment of DLP amino acid sequences from three species from the Dorylaimia (Clade I of Nematoda), *Trichuris muris, Trichinella spiralis* and *Dioctophyme renale*. Also a phylogram from these and other DLP sequences that are not shown - note that the *Soboliphyme baturini* sequence is currently incomplete and not included in the alignment. Multiple alignment and phylogram made with MultAlin set for the Blossum62 matrix. In the consensus line lower case blue font represents amino acids that are identical in two out of the three protein positions, and to illustrate amino acid similarities ! is anyone of IV; $ is anyone of LM; % is anyone of FY; # is anyone of NDQEBZ. The *T. spiralis* sequence was obtained from NCBI accession KAL1244686.1, the *T. muris* from NCBI 6QIX_A; the *D. renale* from NCBI QTE33903.2.

**Figure S6.**
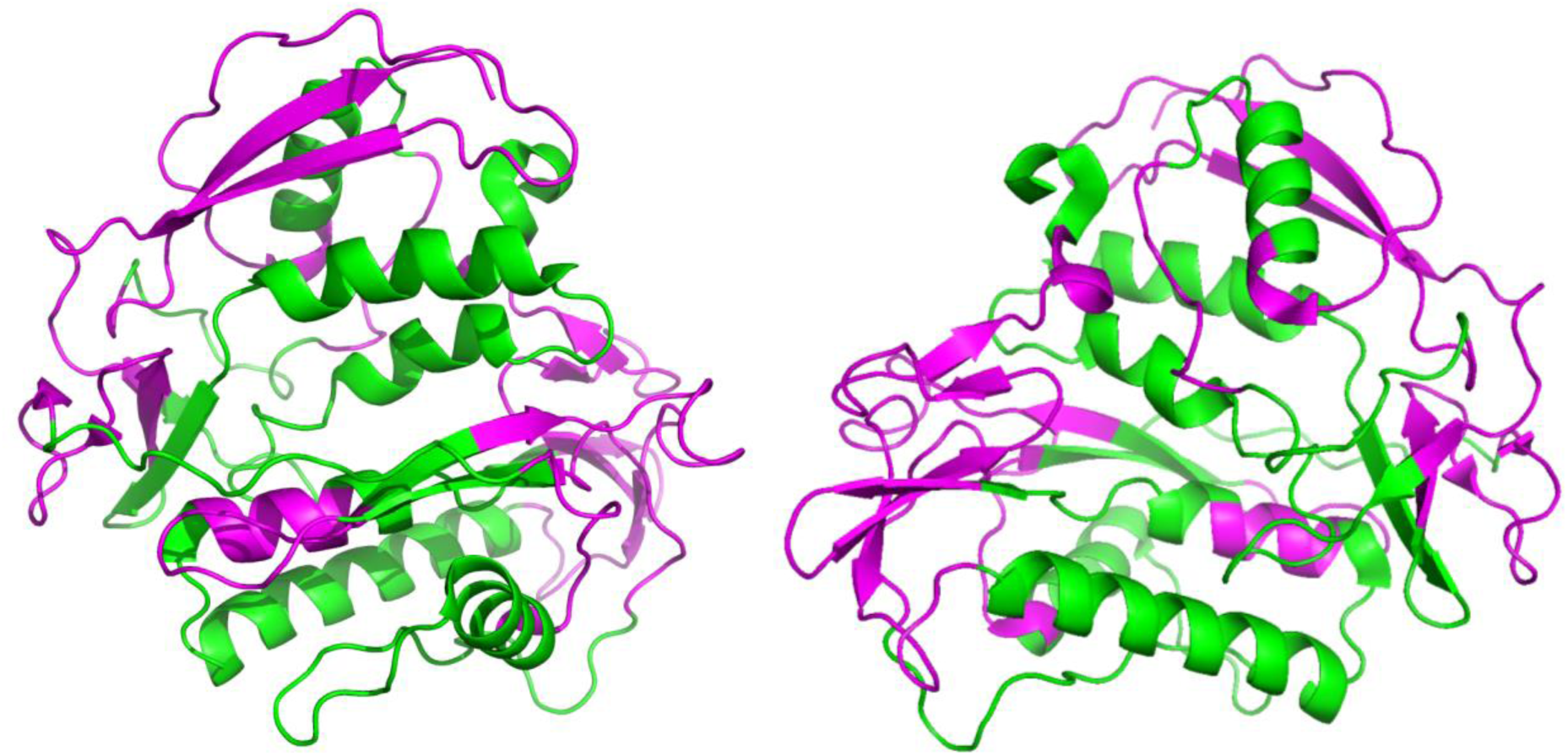
The most conserved regions of the three DLPs aligned in Figure S5 mapped onto p43’s structure. Two orientations of the p43 x-ray crystal structure in which the most conserved regions are coloured magenta as delineated from the amino acids sequence alignments of DLPs from three species of Clade I nematode in Figure S5. These regions were chosen following the rule that each region must have at least four consecutive amino acids that are either identical or biochemically similar (coloured red in the consensus line, and similarity as defined in the legend to Fig 5), and extended in either direction until interrupted by more than two non-identical or biochemically dissimilar amino acids. This shows that the most conserved regions are the extended/β structures and unstructured regions on or near the surface of the protein, potentially therefore involved in interactions with other structures within the nematodes, such as other proteins and membranes, in performing conserved physiological functions.

